# DIY caging apparatus to facilitate chronic and continuous stimulation or recording in an awake rodent

**DOI:** 10.1101/2021.12.16.473031

**Authors:** Syed Faaiz Enam, Brian J. Kang, Johnathan G. Lyon, Ravi V. Bellamkonda

## Abstract

Chronic stimulation of and recording from the brain and brain diseases can require expensive apparatus and tedious cycles of inducing rodents with anesthesia. To resolve this, we have designed and fabricated a low-cost (∼$75 vs. $450) DIY rodent caging apparatus consisting of commercially available and 3D-printed components. This apparatus is customizable and can be used to rapidly prototype devices with large rodent sample sizes. Importantly, it enables continuous and chronic stimulation of and recording from the brains of awake and freely moving rodents. It also opens the possibilities of trying complex paradigms of treatment (continuous, intermittent, variable, and chronic). We have successfully used this caging apparatus for chronic intratumoral hypothermia treatment and are currently using it while advancing electrotactic therapies.

## 1. INTRODUCTION

Work in modern neuroscience and neuroengineering frequently requires stimulation to and recording from the brain through electrodes, optogenetics, temperature-modulation, or pharmaceuticals (1–5). Interventions can be intermittent and chronic (e.g., 2 hours per day for 1 month), but this may be limited by practical considerations including user fatigue and duration of rodent anesthesia. Indeed, in many paradigms, stimulation necessitates inducing the test animal to sleep (6). Although these paradigms can be sufficient, and at times ideal, greater flexibility in stimulation/recording is highly desirable. For example, patterns of treatment that last extended periods of time (e.g., 6 hours/day, overnight, or multiple times per day), are continuous, or complex provide additional therapeutic options to disease conditions. These situations pose two constraints: 1) continuous access to the brain implant; and 2) an awake, alert, and freely moving rodent.

To enable studies that require prolonged or complex treatment/recording paradigms, we have developed a low-cost, customizable caging system for rodents (Fig. 1a). It enables awake and continuous stimulation/recording while allowing freedom of movement through three axes (a slip ring for azimuth rotation, and lever arm for vertical and horizontal displacement). We have previously used this caging system for intratumoral hypothermia treatment (4) and currently for advancing our work with the electrotaxis of brain tumors (7). While caging systems may exist commercially, none are comprehensive, customizable, and low-cost. In comparison, this apparatus is made from 3D-printed and low-cost components. Thus, no sophisticated fabrication equipment is needed; all that is required is access to a 3D printer, soldering iron, and common hardware tools. Adaptations would only require enough CAD expertise to adapt/create 3D printed constructs, and knowledge to choose desired electrical connections. This enables the freedom to replicate and customize the apparatus and eases production requirements to enable studies with large sample sizes.

**Figure 1:**
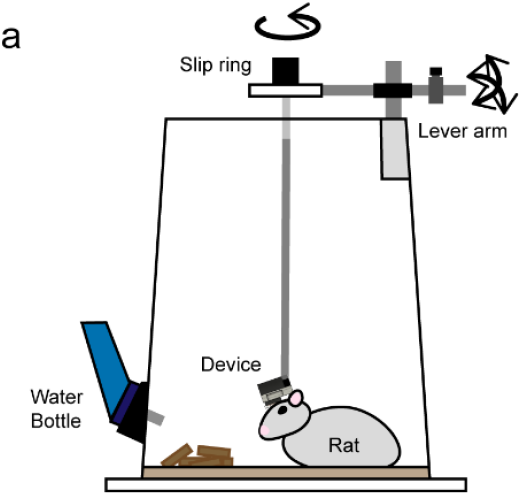
Schematic of a low-cost, customizable caging system for chronic treatment and recording.

## 2. MATERIALS AND METHODS

Materials were purchased from online vendors including McMaster-Carr, Amazon.com, and digi-key. A list of materials with their source and cost is provided (Supplementary Table 1). 3D parts were printed on a MakerGear M3-ID 3D printer with PLA filament (note: most commercially available printers with a single head would be able to complete these prints). Computer-aided drawings for 3D parts were made using AutoDesk Fusion 360 and printed using Octoprint with the CuraEngine slicer. Free design software that are adequate alternatives include the TinkerCAD, FreeCAD, or web-based version of SketchUp.

## 3. RESULTS

### a. Caging to house a rodent

Rodents were housed in a cage which consisted of a platform, inverted bucket, and bottle holder (Fig. 2a). The platform was a 24” x 24” x 1/4” acrylic sheet. The bucket was flipped upside down so that the wider 12” end was centered on the acrylic sheet—more than enough space to house a single rat (8). The height of the bucket was 15” to prevent a rodent from easily escaping. If escape is a concern while the rat is not tethered, a lid or wire mesh can be used as a preventative measure. The height was also important to keep the wiring in the tether cable from bending or winding on itself. The base of the bucket was sawed off (note: sanding with a mill is also possible) to create a hollow tube. Alternatively, a hollow cylindrical pipe of 15” diameter can be used. Plastic bonder was used to adhere the bucket to the platform and set for 24 hours. In the meantime, a bottle holder was 3D-printed (Fig. 2b). When the bond was set, holes were drilled to accommodate the bottle holder and screws (Fig. 2c). To tighten the bottle in the holder, a knob was pressed onto a screw (alternatively, threaded knobs can be purchased).

**Figure 2:**
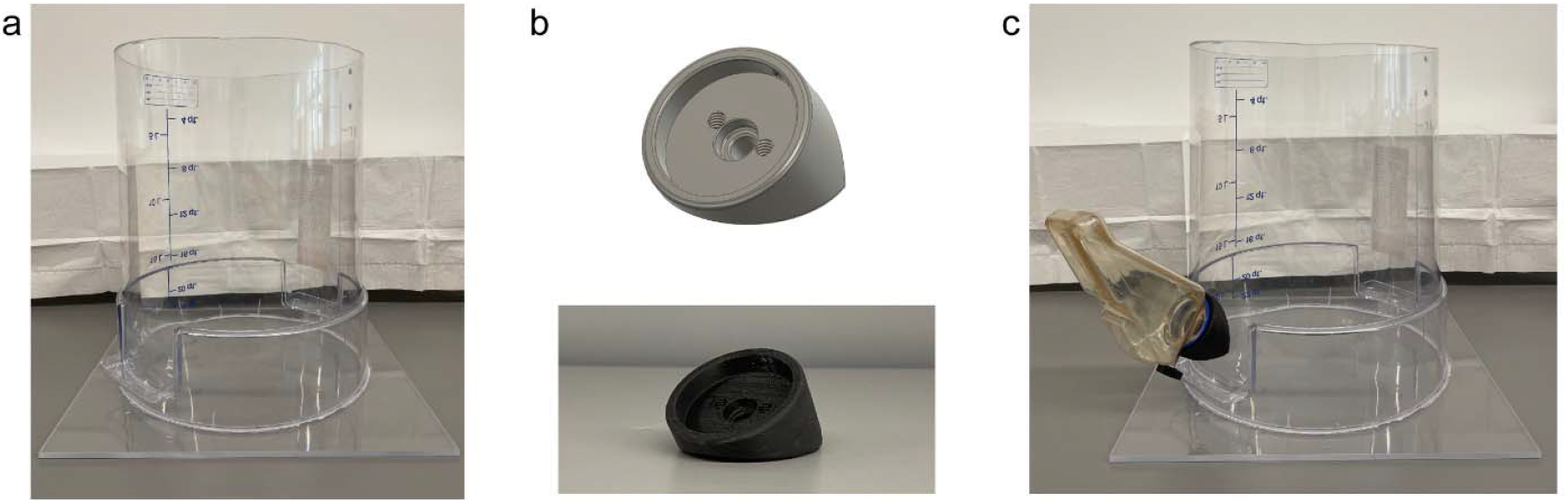
Rodent cage. (a) The cage consists primarily of a cylinder and attached to a large square platform. (b) A water bottle holder designed and used for this cage (above: CAD design, below: 3D print). (c) The cage with a water bottle secured.

### b. Lever arm to enable freedom of movement

A lever arm was developed to provide two axes of movement freedom for both horizontal and vertical displacement. It consisted of four 3D-printed components to provide anchoring and joints and two metal rods. The first 3D-printed part, a receptacle, could be attached to the cage to provide a closed-end socket for a 3/8” metal rod to rotate (Fig. 3a). This provides horizontal movement. Second, two printed parts, to be held together via a partially threaded screw, formed a mobile joint that enabled vertical movement (Fig. 3b). One part of the joint securely attached to the 3/8” rod. The second part of the joint had a hole to pass a ¼” metal rod. At the end of this rod, a 3D-printed platform could be attached to hover above the cage (Fig. 3c). The lever arm consisted of the two joint components and the platform (Fig. 3d). A 100-200g steel counterweight was fabricated and used for balance that could be adjusted along the length of the rod (Fig. 3d). Alternatively, an adjustable 3D part could be designed that holds lead weights. In our setup, the receptacle was attached to the cage with 4 screws and the lever arm was set in the socket (Fig. 3e). For our purposes, the platform held a passive commutator (slip ring) to enable azimuth rotational movement (Fig. 3e). Everything was tightened with screws (threads were included in the 3D parts).

**Figure 3:**
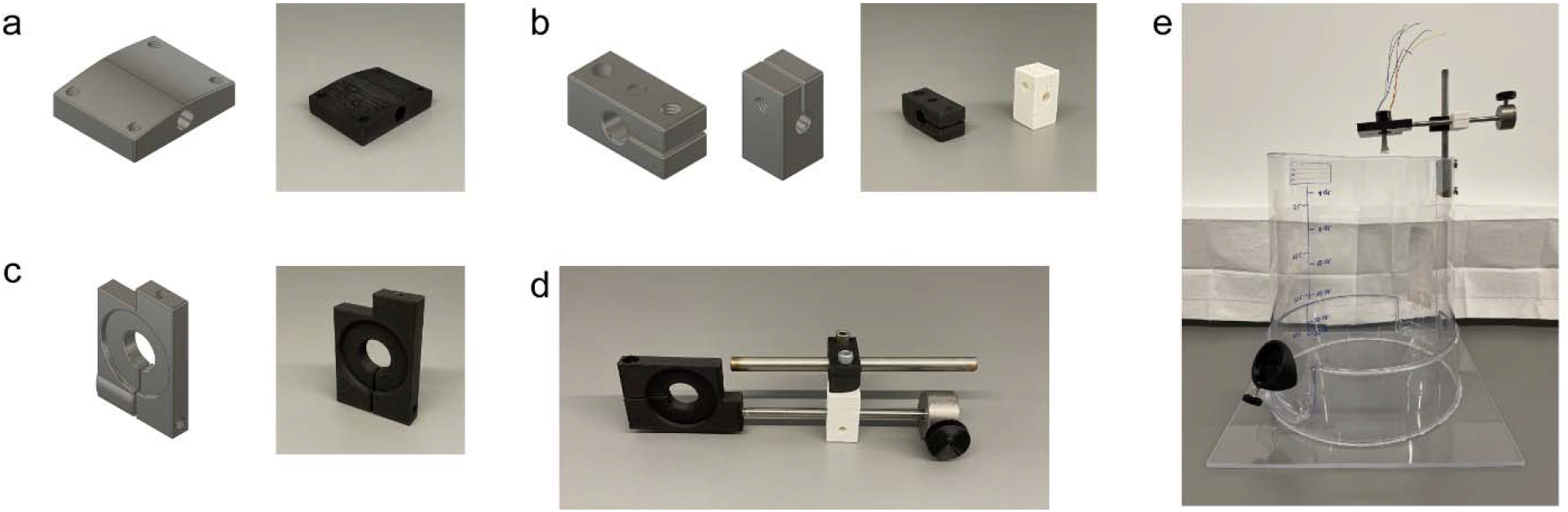
Lever arm. (a) Receptacle for lever arm (left: CAD drawing, right: 3D print). Four threaded holes are visible for attachment to the cage via screws. One, closed-end socket is visible to freely set the 3/8” rod. (b) Joint components (left: CAD drawing, right: 3D prints). With the use of a partially threaded screw, these enable vertical displacement of a ¼” rod. (c) Platform for a slip ring (left: CAD drawing, right: 3D print). This part attaches to a ¼” rod and provides a surface to secure a slip ring. The hold is tightened with a screw. (d) The lever arm on its side with its rods and counterweight. (e) The lever arm with a slip ring sitting in the receptacle and hovering over the cage. The counterweight can be adjusted to provide balance.

### c. Wiring and external system for treatment and recording

Apart from the structural components of the cage, for our applications (4,7) we required electronic components to be powered and recorded from. To enable this, the cable that tethered the animal to our commutator comprised of 28G jumper wires with male connectors on either end enclosed in a spring sheath (Fig. 4a). To secure the connection, medical-grade epoxy was applied at the joints (cured for 1-2 days) and further supported with hot glue (Fig. 4b). The surrounding spring was used to relieve tension on the wires and joints and provide an extra layer of protection from animal contact (Fig. 4c). For further strength, a layer of epoxy was added on top of the dried glue. It was important to be careful during the epoxy and glue steps to not let any glue seep into the connector (this may make the connectors unattachable). Alternatively, the spring can be secured on one end only to the male connector. In that case, a ring of hot glue was added around the spring end to prevent damaging the wire. The cables were the most frequent point of failure and thus making extra is convenient (substituting higher gauge wires may also provide more strength). Additionally, when soldering the circuit, it is preferable to have the outer wires carry sensory/recording information as the outer ones tend to break first.

**Figure 4:**
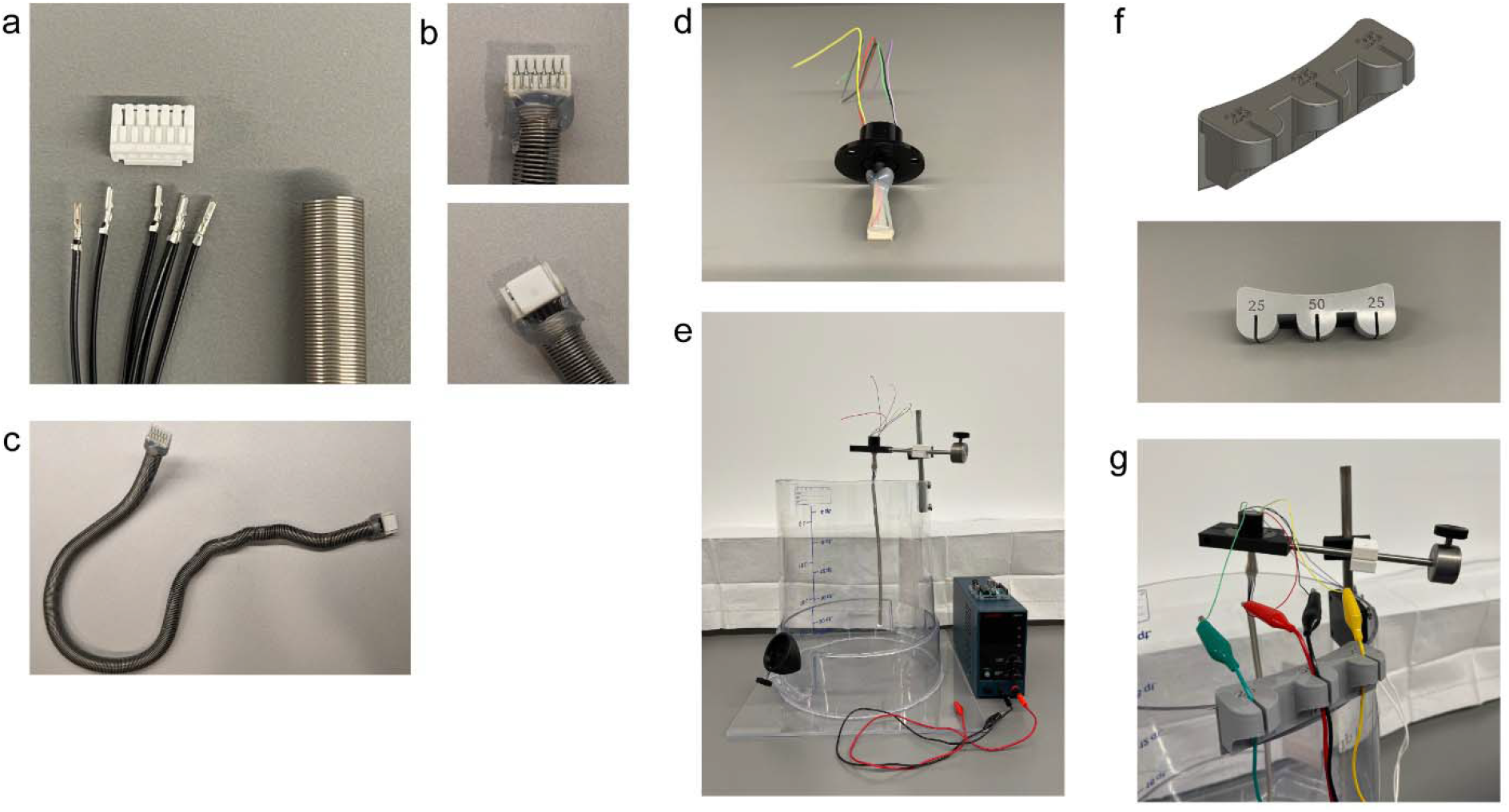
Electrical wiring system. (a) Picture of the components of one end of the wiring tether used in our setup. This includes a male connector (white), jumper wires (available crimped), and a spring to protect the wires. (b) The connector, wire, and spring assembled with epoxy and hot glue to secure the joint. (c) The complete tether/cable with the spring stretched from one end to the other. (d) Slip ring with wires soldered to a female connector and protected with epoxy (at the joint) and hot glue all around. (e) Example of a cage with wiring and external equipment on the side. (f) A component to manage connecting cables (above: CAD drawing, bottom: 3D print). (g) Picture demonstrating the cable holder hanging from the side of the cage and securing cables with alligator clips to connect external devices to the slip ring.

The slip-ring wires were soldered to a female connector and then protected by epoxy and hot-glue (Fig. 4d). Outside the cage, we used either a power supply or function generator for stimulating, and Arduino or oscilloscope for recording (Fig. 4e). Cables from these devices were connected to end wires of the slip ring via alligator clips. To manage the cables, a holder was 3D printed to hang on the sides of the cage (Figs. 4f and g). While tape can also be used, this holder makes changing cages and connecting/disconnecting cables easier.

### d. Use of housing apparatus and maintenance

Once prepared, the cages were washed with water and then cleaned with veterinary disinfectant (e.g., RESCUE™). After wiping dry, bedding and nest material were added, rat food was placed on the floor, water bottles were filled, and a wooden block was provided for enrichment. Rats were singly housed and given a week to accustom to the cage prior to beginning any procedures (Fig. 5a). Afterwards, they were maintained and treated in their cage (Fig. 5b and c) for up to 3 months. The cages were cleaned 1-2 times per week by dumping out the bedding, washing thrice with a pressure hose, and then sanitizing with veterinary disinfectant. Because of the relative low-cost of this caging, we had extra cages on hand that made it easy to maintain both the cages and the treatment of the rats. When a cage needed to be swapped, the rat with the tether and lever arm could be quickly placed into a freshly prepared cage with minimal disruption to stimulation/recording. Altogether, this system provided ease to conduct this experiment in up to numerous rats simultaneously (4 rats shown in Fig. 5d but up to 18 singly housed rats have been tested in one experiment) with a central computer to record data (Fig. 5d).

**Figure 5:**
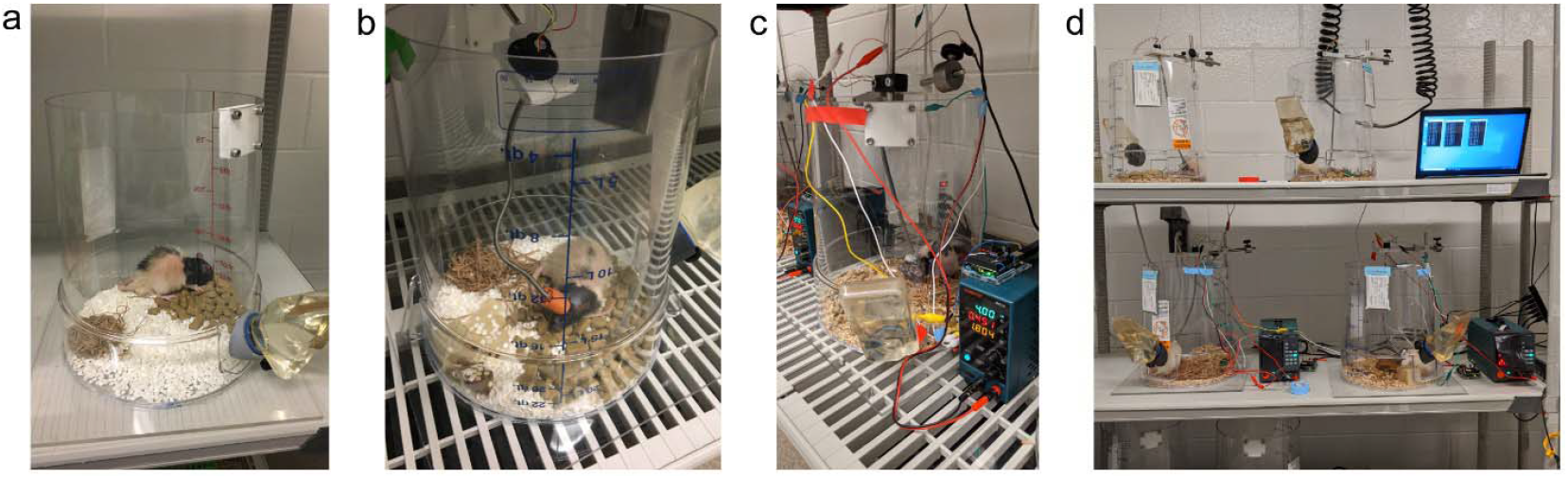
Rats under protocol in the custom cage. (a) A rat becoming accustomed to the cage with fresh bedding, nest, water, and food. (b) A rat with a brain tumor undergoing treatment and recording of electrical stimulation. (c) A rat with a brain tumor under treatment with a powered hypothermia device. Here, temperature is recorded through the visible Arduino. (d) A small set up of 4 rat cages with treatment systems and a central computer for temperature recording.

## 4. DISCUSSION

Here we provided designs for a low-cost, customizable caging setup to enable continuous and chronic stimulation and recording of awake rodents. The apparatus allows the rats to move freely while undergoing treatment or recording. The 3D printed components are customizable as needed for specific situations. Because of the cost of the parts, cages can be rapidly prototyped and produced in scale for *in vivo* experiments with larger sample sizes. The average cost for one cage (with lever-arm and slip ring) was ∼$75 (Supplementary Table 1). A non-customizable but commercially available cage (with a lever-arm and swivel for fluid delivery) would cost ∼$450 (9). The accessibility to this apparatus opens therapeutic treatment paradigms utilizing both complex patterns and targeting complex diseases.

One disease that might particularly benefit from prolonged or complex treatments is a brain tumor. For example, we have previously demonstrated that continuous or intermittent (18 hours/day) of local cytostatic hypothermia can halt the growth of glioblastoma *in vitro* (4). In this set of studies, we used the above-described apparatus for continuous hypothermia delivery *in vivo*. We have also observed that the application of electric fields can promote tumor migration (7). However, moving tumor cells from one location to another would necessitate continuous or prolonged stimulation periods/regimens. Thus, we are currently advancing this work with the use of this apparatus. Outside of our lab, other labs have demonstrated that, for example, neural activity promotes glioma proliferation—this apparatus could be used to assess prolonged and complex inhibitory stimulation patterns (10). While these treatment paradigms would be ripe for cancer treatment, they can also be used for dementias, stroke, and brain injury (2). Beyond the brain, this apparatus could be modified to work with peripheral nerve electrodes and stimulation (11) as well as any other wired electronic system outside the cranium.

Some applications may require minor or major modifications, all of which can be readily adapted from our design. For example, for optogenetics, an optical commutator can be used in place of the current passive slip ring we used. This would enable continuous, complex, and chronic optogenetic stimulation patterns. For concurrent drug treatment, a combined electric and fluidic commutator or slip ring can be purchased or prototyped. Alternatively, for experiments that require more precision/less noise, an active commutator can be fit into the apparatus. To monitor rodent behavior, a webcam holder can be printed and hung by the side of the cage. The tethered cable could be used to provide power or a bidirectional electrical signal. Control circuity could be outboard or local with the implant. Thus, this system could be used for virtually any experimental system where a wired electrical setup is required.

To treat chronic diseases with continuous or complex treatment paradigms, or recording for long periods, caging is needed that enables an awake rodent to behave freely. While advances in wireless systems are underway, those require sophisticated redesigns and can be challenging in the prototyping stage (or for experiments requiring large amounts of power). Alternatively, these cages are low-cost and highly tailorable to specific needs and may enable opportunities for future scientists to test/record continuous and complex paradigms.

---

**Supplementary Table 1:**
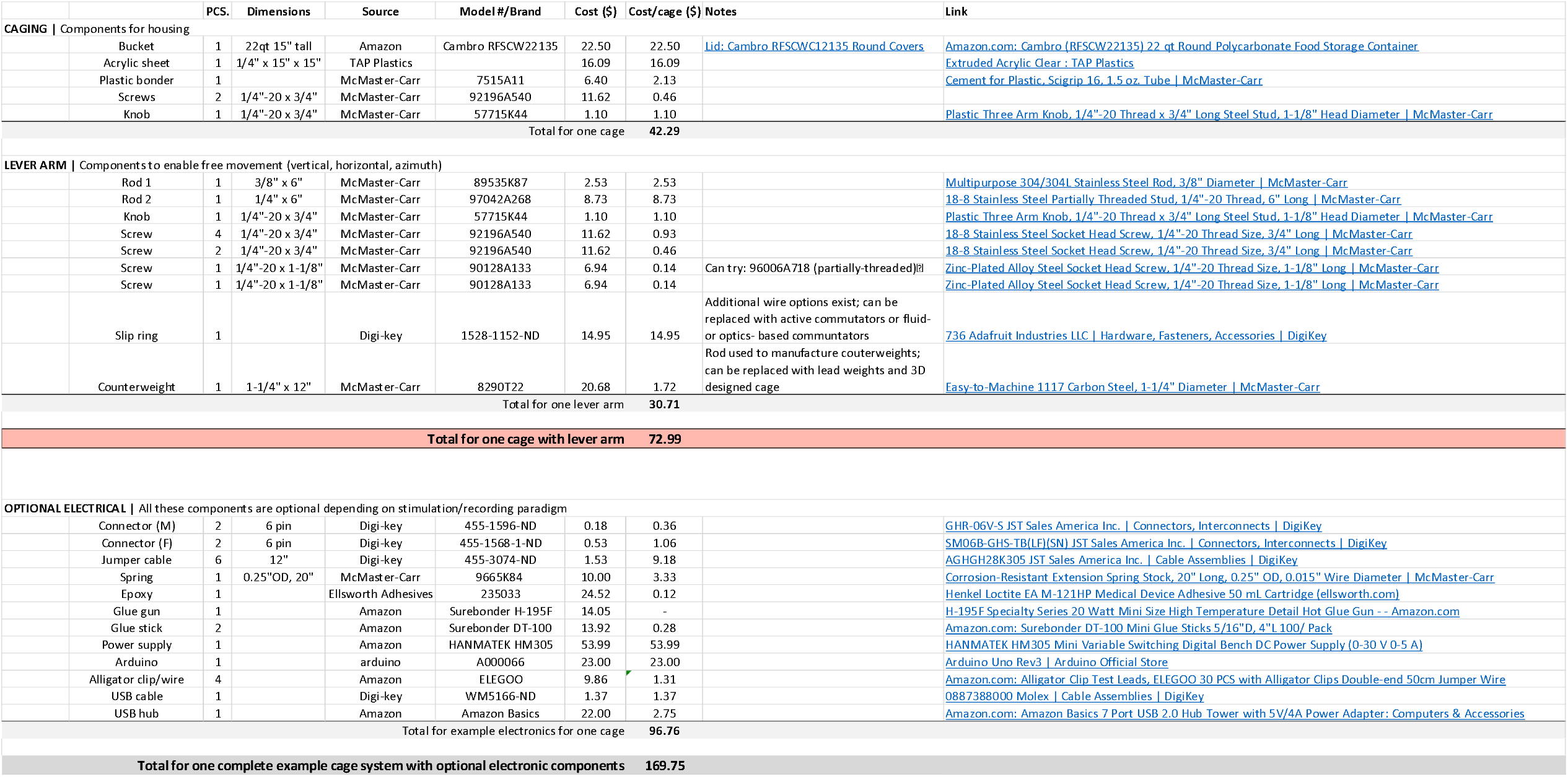
Components and their source and cost for DIY caging apparatus separated into categories. The caging described requires most components from the “Caging” and “Lever Arm” sections. The slip ring and counterweight can be replaced as needed. Components in the “Optional Electrical” section are dependent on the target application.

